# A Robust Combination Test Using the Normal Distribution, with Application to Phylogenetic Inference

**DOI:** 10.1101/2023.07.09.548262

**Authors:** Md Rejuan Haque, Laura Kubatko

## Abstract

Combination tests are used to combine P-values from individual studies to test a global null hypothesis. These types of tests can also be applied to combine P-values from testing separate null hypotheses within the same study in cases for which the procedure for testing a global null hypothesis is unavailable. One such application of a combination test is to detect the presence of hybrid species within a set of species. Although many combination tests have been proposed in the literature, there is no uniformly most powerful test applicable for all conditions. For instance, in the hybrid detection application, it is expected that only a few of the species within a set species might have truly arisen via hybridization, and thus when tested, only a few of the individual P-values are expected to be significant. Thus, a desirable property of a combination test for this situation is to be able to reject the global null hypothesis even if only a small fraction of the individual P-values are significant. In this paper, we propose a new combination test based on the normal distribution that assigns weight to each test adaptively, thereby providing powerful results even when only a small fraction of the individual null hypotheses are false. A comprehensive simulation study, as well as real data applications, demonstrate that the proposed test is powerful in detecting false global null hypotheses under several situations for which existing tests have low power.

## 1 Introduction

Combination tests are used to combine the results of individual hypothesis tests, often arising from independent studies, to draw reliable and generalizable conclusions about an underlying global null hypothesis. By pooling data from multiple studies, statistical combination tests can provide more accurate estimates of the effect size and increase statistical power beyond what may be possible with individual studies alone. Additionally, these tests can help identify sources of heterogeneity or inconsistency across studies and provide insight into the factors that may influence the relationships between variables of interest. Other statistical approaches, such as meta-analysis, aim to combine results from multiple independent studies. However, unlike a combination test, meta-analysis requires several statistics, such as 95% confidence intervals or odds ratios (Chen, Yang, Liu, Yang, Li and Yang, 2014), which may not be available for all individual studies. Instead, we might only have access to the P-values from each study, and we may thus be unable to use meta-analysis. In this case, a combination test can be used to combine the individual P-values to test a global null hypothesis.

A combination test works by aggregating information from individual tests within a single study to perform a comprehensive global test (Liu and Xie, 2020). This kind of test can be used to evaluate the global null hypothesis that none of the individual null hypotheses are false against the alternative that at least one is false. Combination tests are particularly beneficial when executing a global test is difficult or impossible but individual tests are easy to carry out. An instance of a useful application of combination tests is in detecting the presence of hybrid species (species that share their genetic ancestry with two distinct parental species) among a set of species. While methods for testing whether a given species is hybrid or not when only four species are considered at a time exist (Haque and Kubatko, 2023; Kong and Kubatko, 2021; Kubatko and Chifman, 2019; Meng and Kubatko, 2009), no methods have been proposed to detect whether or not a data set with more than four species contain one or more hybrid species. This problem is important because the evolutionary relationships among a set of species with one or more hybrid species is depicted using a species network, whereas a species tree is used for data without hybrid species. Hence, a global test for the presence of hybrid species should precede any phylogenetic analysis intended to estimate the evolutionary relationships among a set of species. Since no methods are available to perform a global test of hybridization, one can use a combination test to combine the P-values of individual tests that consider four species at a time. However, when choosing a proper combination test for this scenario, one should keep in mind that only a small fraction of the individual tests are expected to be significant as hybrid speciation is rare, and thus it is unlikely that a data set will contain a large number of hybrids. Also, the testing procedure is further complicated by gene-tree species-tree discordance in phylogenetics (Wicke et al., 2023).

The statistical literature is replete with methods for combining P-values to test a global null hypothesis (Birnbaum, 1954; Fisher, 1992; Good, 1955; Lancaster, 1961; Lipták, 1958; Mosteller and Bush, 1954; Pearson, 1938; Stouffer et al., 1949; Tippett et al., 1931; Whitlock, 2005). Despite the availability of numerous methods, no single combination test has been shown to be uniformly most powerful for all situations (Birnbaum, 1954; Krishnamoorthy et al., 2022). For instance, in the hybridization detection studies mentioned above, the rarity of hybrids often leads to a small number of significant P-values and a large number of nonsignificant P-values. Moreover, in the absence of hybrid species, hybrid detection tests often produce P-values close to 1 (Blischak et al., 2018; Kubatko and Chifman, 2019), which results in large nonsignificant P-values in the combination test, thereby reducing the power. Therefore, in this paper, we propose a new combination test based on the normal distribution that has substantive power even when only a small fraction of the individual null hypotheses are false and a large number of the individual tests give P-values close to 1. Our test incorporates the sample sizes in individual studies, leading to a robust test that is applicable in many situations. In section 2, we discuss some existing popular combination tests. In section 3, we discuss our proposed test. Section 4 gives the results of the application of our test to simulated data, as well as to several empirical data sets. In section 5, we discuss the advantages of our test as well as some limitations.

## 2 Overview of Existing Combination Tests

Let *P*_1_, *P*_2_, *· · ·, P*_*j*_, *· · ·, P*_*k*_ be P-values arising from testing *k* independent null hypotheses *H*_0*j*_ vs. alternative hypotheses *H*_1*j*_; *j* = 1, 2, *· · ·, k*. We assume that under each individual null hypothesis, the P-value (*P*_*j*_) follows a Uniform distribution on the interval (0, 1), i.e., *P*_*j*_ ∼ Uniform(0, 1). Now, consider testing the following global null hypothesis:

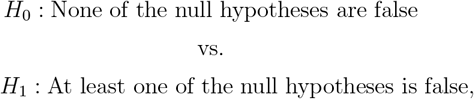

i.e., *H*_0_: ∩*H*_*j*0_ vs. *H*_1_: ∪*H*_*j*1_; *j* = 1, 2, · · ·, *k*. One way to perform this global test is by using a combination test. Several popular combination tests have been proposed in the statistical literature. We review several of these below.

### 2.1 Fisher’s Combination Test

Fisher (1992) proposed the oldest and arguably most powerful combination test for many situations. Studies have demonstrated that Fisher’s test is particularly useful for analyzing genetic data targeted to uncover genes associated with a particular disease (Chen, Huang and Liu, 2014; Chen et al., 2012; Chen, Huang and Ng, 2014; Chen et al., 2016; Chen and Ng, 2012), and it yields robust and powerful results. The test statistic for Fisher’s combination test is: 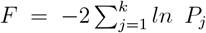. Since *P*_*j*_ ∼ Uniform(0, 1) under *H*_0*j*_, *−*2 *ln P*_*j*_ has a 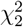 distribution, and consequently, the statistic *F* has a 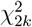 distribution under the global null hypothesis *H*_0_. Let *p*_1_, *p*_2_, · · ·, *p*_*k*_ be the observed P-values from the k individual tests, and let 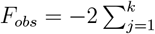 *ln p*_*j*_ be the observed value of the test statistic *F*. To test the global null hypothesis *H*_0_ at a given significance level *α*, one should reject *H*_0_ if

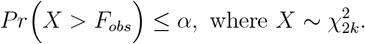

### 2.2 The Inverse Normal Test

The Inverse Normal test (Z-test), proposed by Stouffer et al. (1949), is based on the normal distribution. Though Fisher’s test performs well in diverse scenarios, depending on the effect sizes and directions of individual tests, the performance of the Inverse Normal test may surpass that of Fisher’s test. Specifically, if all the effect sizes are of similar magnitude and direction, the Z-test performs better than Fisher’s test (Chen, Yang, Liu, Yang, Li and Yang, 2014). The test statistic for the Inverse Normal test is:

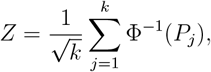

where Φ(*·*) is the cumulative distribution function of the standard normal distribution. Because each *P*_*j*_ is assumed to have an independent Uniform(0, 1) distribution under the individual null hypotheses *H*_0*j*_, Φ^−1^(*P*_*j*_) will have independent standard normal distributions, and hence the statistic *Z* will have a standard normal distribution under the global null hypothesis *H*_0_. Thus, if the observed P-values are *p*_1_, *p*_2_, · · ·, *p*_*k*_, and the observed test statistic is 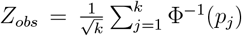, given a significance level *α*, one should reject the global null hypothesis *H*_0_ if *Z*_*obs*_ ≤ *z*_*α*_, where *z*_*α*_ is the 100*α* percentile of the standard normal distribution.

### 2.3 The Weighted Inverse Normal Test

The Inverse Normal test assumes equal weight for each of the individual tests. However, to improve power, some researchers have incorporated sample sizes and standard errors of each test from the individual studies into the Inverse Normal test by introducing a weight component, leading to the Weighted Inverse Normal test (Stouffer et al., 1949; Whitlock, 2005). Whitlock (2005) argued that assigning more weight to the tests with larger sample sizes might be beneficial and proposed the following test statistic

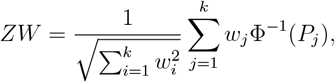

where *w*_*i*_ = *n*_*i*_ *−* 1 and *n*_*i*_ is the sample size of the *i*^*th*^ individual test. Under the global null hypothesis *H*_0_, the test statistic *ZW* follows the standard normal distribution. Thus, if the observed P-values are *p*_1_, *p*_2_, *· · ·, p*_*k*_, and the observed test statistic is 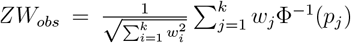, given a significance level *α*, one should reject the global null hypothesis *H*_0_ if *ZW*_*obs*_ ≤ *z*_*α*_, where *z*_*α*_ is the 100*α* percentile of the standard normal distribution.

### 2.4 The Inverse χ^2^ Test

Methods based on the chi-squared distribution have also been proposed (Krishnamoorthy et al., 2022; Lancaster, 1961). For instance, Lancester (1961) converted the P-values from the separate studies into a chi-square variate and added them to get a test statistic for a global test. Lancester’s test is a generalization of Fisher’s test and is the same when the degrees of freedom are set to two for each study. Recently, Krishnamoorthy et al. (2022) proposed the Inverse χ^2^ test, which generalizes Lancester’s test by assigning more weight to individual tests with larger sample sizes. The test statistic proposed by Krishnamoorthy et al. (2022) is:

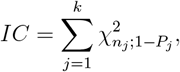

where 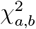 is the 100*b*^*th*^ percentile of the chi-square distribution with df *a*. Thus, under the global null hypothesis *H*_0_, the test statistic *IC* follows a 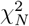 distribution, where 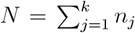. Hence, if the observed P-values are *p*_1_, *p*_2_, *· · ·, p*_*k*_, and the observed test statistic is 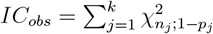. Given a significance level *α*, one should reject the global null hypothesis *H*_0_ if 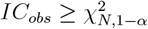.

## 3 Methods

### 3.1 Random Inverse Normal Test: RINT

Combination tests require appropriate weighting to maximize power if the global null hypothesis fails to explain the generated data. Importance should also be given to the sample size of each of the individual tests, as a large sample size might provide more information about *H*_0_. We propose a test statistic based on the normal distribution that defines weights for each test adaptively.

Consider combining P-values *P*_1_, *P*_2_, *· · ·, P*_*k*_ from *k* individual tests each with sample size *n*_1_, *n*_2_, *· · ·, n*_*k*_ to test the global null hypothesis *H*_0_. We propose the following test statistic:

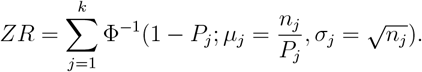

Notice that the *j*^*th*^ individual P-value *P*_*j*_ is converted to the (1 − *P*_*j*_)100 percentile of a normal distribution with mean *μ*_*j*_ = *n*_*j*_*/P*_*j*_ and standard deviation 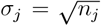. The intuition is that when *P*_*j*_ is small, *μ*_*j*_ is large, leading to a large value of the test statistic. Also, tests with larger sample sizes will lead to a larger mean and a larger standard deviation, which in turn, will result in a larger value of the test statistic since 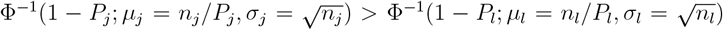 when *P*_*j*_ = *P*_*l*_ and *n*_*j*_ *> n*_*l*_. Hence, the proposed test assigns larger weights to the individual tests with smaller P-values and/or larger sample sizes.

One issue with the proposed statistic is that it does not have any known distribution under the global null hypothesis, as the mean and the standard deviation corresponding to each individual P-value are random. However, under each of the individual null hypotheses *H*_0*j*_, we assume that *P*_1_, *P*_2_, *· · ·, P*_*k*_ are Uniform(0, 1) random variable, and using this assumption, we can generate data from the distribution under the global null hypothesis *H*_0_ in order to simulate the null distribution. To test the global null hypothesis *H*_0_, one can follow the following steps:

1. For observed individual P-values *p*_1_, *p*_2_, *· · ·, p*_*k*_ with sample sizes *n*_1_, *n*_2_, *· · ·, n*_*k*_, compute the observed value (*ZR*_*obs*_) of the statistic *ZR*.
2. Generate *k* values (*x*_1_, *x*_2_, *· · ·, x*_*k*_) from the Uniform(0, 1) distribution.
3. Using *x*_1_, *x*_2_, *· · ·, x*_*k*_ as the P-values and *n*_1_, *n*_2_, *· · ·, n*_*k*_ as the sample sizes, calculate the value of the statistic *ZR* and call it *ZR*_01_.
4. Repeat steps 2-3 *B* times (e.g., *B* = 10^8^) to generate *B* observations from the distribution of the test statistic under the global null hypothesis *H*_0_. Let the generated values be *ZR*_01_, *ZR*_02_, *· · ·, ZR*_0*B*_.
5. Calculate the P-value: 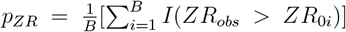, where *I*(*·*) is the indicator function.
6. Reject the global null hypothesis *H*_0_ with significance level *α* if *p*_*ZR*_ *≤ α*.

### 3.2 Simulation Study

To evaluate the performance of the proposed test, we carried out an extensive simulation study that examined various scenarios. These scenarios involved different sample sizes for the individual tests, different numbers of individual tests, and different standard deviations used to generate the individual test statistics. This approach allowed us to thoroughly analyze the performance of the proposed test under diverse conditions and determine its statistical properties, such as power and Type I error rate. The general data generation from an individual test (*H*_0*i*_: *μ*_*i*_ = 0 *vs. H*_1*i*_: *μ*_*i*_ *>* 0) when for instance, *μ*_*i*_ = *μ, n*_*i*_ = *n*, and *σ*_*i*_ = *σ* has the following three-step process:

- Step 1: Generate a value of 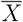 using the normal distribution as follows:

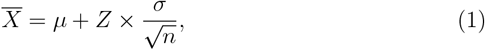

where *Z ∼* Normal(0, 1).
- Step 2: Generate the value of sample variance (*S*^2^) using the chi-squared distribution as follows:

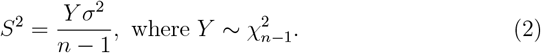
- Step 3: Compute P-values for each individual null hypothesis testing *H*_0*j*_: *μ* = 0 using the standard one-sample t-test.

This procedure is repeated *k* times to generate *k* P-values. We allow the values of *n* and *σ* to vary across replicates as follows: *n* ∼ Poisson(*λ*); *σ* ∼ Uniform(*a, b*). We also allowed fixed values for *n* and *σ*. We also consider various choices for the number of individual null hypotheses (*k*) to be combined. We describe the various choices of *λ, a*, and *b* below. We also consider different scenarios for which the number of true null hypotheses varies, and we consider different values of *μ* under the alternative.

To evaluate the performance of the proposed test, we considered the following 11 scenarios and repeated each scenario for *α* = 0.05 and *α* = 0.01. We also repeated each of the scenarios using *S*^2^ generated from Uniform(1, 10). This choice of *S*^2^ makes it possible for individual tests to vary substantially and makes it hard to reject the null hypothesis. All the simulation scenarios were repeated 1000 times and the power of the proposed combination test along with several other existing tests was calculated.

- Scenario 1: P-values were generated for two individual studies testing *H*_0*i*_: *μ*_*i*_ *≤* 0 *vs. H*_1*i*_: *μ*_1*i*_ *>* 0; *i* = 1, 2. The sample sizes for each study were generated from a Poisson distribution with a mean of 50. Standard deviations *σ*_*i*_ for each study were generated from the Uniform(2,4) distribution. Statistics under the individual null hypotheses were generated using *μ* = 0, and statistics under the alternative hypotheses were generated using 0 *< μ ≤* 3.
- Scenario 2: P-values were generated for two individual studies testing *H*_0*i*_: *μ*_*i*_ *≤* 0 *vs. H*_1*i*_: *μ*_1*i*_ *>* 0; *i* = 1, 2. The sample sizes for each study were generated from a Poisson distribution with a mean of 50. The standard deviations *σ*_*i*_ for each study were generated from the Uniform(2,4) distribution. Statistics under the individual null hypotheses were generated using *μ* = − 2, making P-values under the null hypotheses close to 1, and statistics under the alternative hypotheses were generated using 0 *< μ ≤* 3.
- Scenario 3: P-values were generated for ten individual studies testing *H*_0*i*_: *μ*_*i*_ ≤ 0 *vs. H*_1*i*_: *μ*_1*i*_ *>* 0; *i* = 1, 2, *· · ·*, 10. The sample sizes for each study were generated from a Poisson distribution with a mean of 50. The standard deviations *σ*_*i*_ for each study were generated from the Uniform(2,4) distribution. Statistics under the individual null hypotheses were generated using *μ* = − 2, making P-values under the null hypothesis close to 1, and statistics under the alternative hypotheses were generated using 0 *< μ ≤* 3.
- Scenario 4: P-values were generated for twenty individual studies testing *H*_0*i*_: *μ*_*i*_ ≤ 0 *vs. H*_1*i*_: *μ*_1*i*_ *>* 0; *i* = 1, 2, *· · ·*, 20. The sample sizes for each study were generated from a Poisson distribution with a mean of 50. The standard deviations *σ*_*i*_ for each study were generated from the Uniform(2,4) distribution. Statistics under the individual null hypotheses were generated using *μ* = 0, and statistics under the alternative hypotheses were generated using 0 *< μ ≤* 3.
- Scenario 5: P-values were generated for twenty individual studies testing *H*_0*i*_: *μ*_*i*_ ≤ 0 *vs. H*_1*i*_: *μ*_1*i*_ *>* 0; *i* = 1, 2, *· · ·*, 20. The sample sizes for each study were generated from a Poisson distribution with a mean of 30. The standard deviations *σ*_*i*_ for each study have been generated from the Uniform(2,4) distribution. Statistics under the individual null hypotheses were generated using *μ* = 0, and statistics under the alternative hypotheses were generated using 0 *< μ ≤* 3.
- Scenario 6: P-values were generated for twenty individual studies testing *H*_0*i*_: *μ*_*i*_ ≤ 0 *vs. H*_1*i*_: *μ*_1*i*_ *>* 0; *i* = 1, 2, *· · ·*, 20. The sample sizes for each study were generated from a Poisson distribution with a mean of 100. The standard deviations *σ*_*i*_ for each study were generated from the Uniform(2,4) distribution. Statistics under the individual null hypotheses were generated using *μ* = 0, and statistics under the alternative hypotheses were generated using 0 *< μ ≤* 3.
- Scenario 7: P-values were generated for twenty individual studies testing *H*_0*i*_: *μ*_*i*_ ≤ 0 *vs. H*_1*i*_: *μ*_1*i*_ *>* 0; *i* = 1, 2, · · ·, 20. The sample sizes for each study were generated from a Poisson distribution with a mean of 50 but sorted in a way so that test statistics under the null parameter space have smaller sample sizes compared to those under the alternative parameter space. The standard deviations *σ*_*i*_ for each study were generated from the Uniform(2,4) distribution. Statistics under the individual null hypotheses were generated using *μ* = 0, and statistics under the alternative hypotheses were generated using 0 *< μ ≤* 3.
- Scenario 8: P-values were generated for twenty individual studies testing *H*_0*i*_: *μ*_*i*_ ≤ 0 *vs. H*_1*i*_: *μ*_1*i*_ *>* 0; *i* = 1, 2, · · ·, 20. The sample sizes for each study were generated from a Poisson distribution with a mean of 50 but sorted in a way so that test statistics under the null parameter space have bigger sample sizes compared to those under the alternative parameter space. The standard deviations *σ*_*i*_ for each study were generated from the Uniform(2,4) distribution. Statistics under the individual null hypotheses were generated using *μ* = 0.
- Scenario 9: P-values were generated for twenty individual studies testing *H*_0*i*_: *μ*_*i*_ *≤* 0 *vs. H*_1*i*_: *μ*_1*i*_ *>* 0; *i* = 1, 2, *· · ·*, 20. The sample sizes for each study were fixed at 70. The standard deviations *σ*_*i*_ for each study were fixed at 2. Statistics under the individual null hypotheses were generated using *μ* = 0, and statistics under the alternative hypotheses were generated using 0 *< μ ≤* 3.
- Scenario 10: P-values were generated for one hundred individual studies testing *H*_0*i*_: *μ*_*i*_ ≤ 0 *vs. H*_1*i*_: *μ*_1*i*_ *>* 0; *i* = 1, 2, · · ·, 100. The sample sizes for each study were generated from a Poisson distribution with a mean of 50. The standard deviations *σ*_*i*_ for each study were generated from the Uniform(2,4) distribution. Fifty statistics under the individual null hypotheses were generated using *μ* = − 2, making the corresponding P-values under the null hypothesis close to 1. The rest of the statistics under the null hypotheses were generated using *μ* = 0, and statistics under the alternative hypotheses were generated using 0 *< μ ≤* 3.
- Scenario 11: P-values were generated for twenty individual studies testing *H*_0*i*_: *μ*_*i*_ = 0 *vs. H*_1*i*_: *μ*_1*i*_ ≠ 0; *i* = 1, 2, · · ·, 20. The sample sizes for each study were generated from a Poisson distribution with a mean of 50. The standard deviations *σ*_*i*_ for each study were generated from the Uniform(2,4) distribution. Statistics under the individual null hypotheses were generated using *μ* = 0, and statistics under the alternative hypotheses were generated using *−*2 *≤ μ <* 0 and 0 *< μ ≤* 2.

## 4 Results

### 4.1 Simulation Studies

We designed 11 different simulation scenarios to assess the performance of the proposed test (RINT) compared to several other existing tests. We consider changing the sample size, standard deviation, number of individual tests, and value of the mean (*μ*) under the null and the alternative parameter spaces used to generate the individual P-values. Panels a and b of Figure 1 illustrate the power of each method when combining only two individual tests. In panel a, the test statistics under the null distribution were generated using *μ* = 0 (Scenario 1), while panel b displays results generated with *μ* = − 2 (Scenario 2). This leads to a larger P-value under the true null hypothesis, making it closer to 1. In the first column, only one of the null hypotheses is true (and thus the number of true alternatives, N.alt, is 1). In the second column, neither null hypothesis is true (i.e., N.alt = 2). Panel a of Figure 1 shows that when one of the two individual nulls is true, the proposed test has similar power to Fisher’s test. When both null hypotheses are false, all tests perform similarly. However, panel b shows that if one of the individual nulls is true, producing a P-value closer to 1 with the other P-value expected to be smaller than the significance level (in this case, 0.05), the proposed test outperforms all the other tests. It is noteworthy that when both P-values are expected to be smaller than the significance level, the proposed test has slightly less power than the other tests.

**Figure 1:**
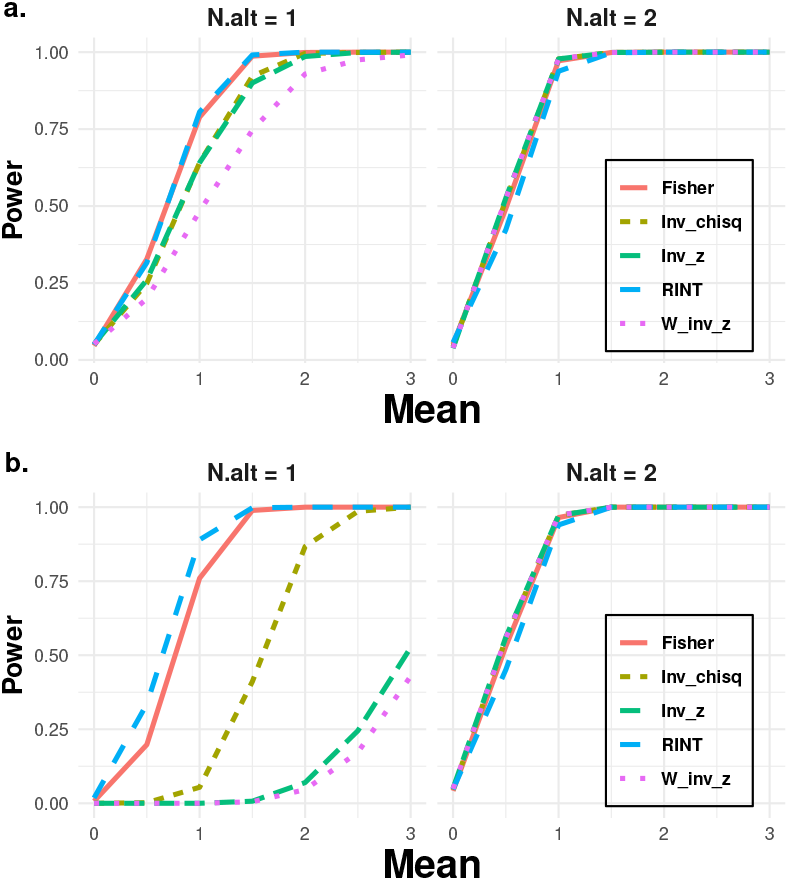
Plots of the power of the combination tests for data sets for which the P-values were generated under scenario 1 (panel a) and scenario 2 (panel b) with *α* = 0.05. The x-axis of the plot shows the value of *μ* under the alternative space used to generate the individual test statistic, and the y-axis shows the power of several combination tests. P-values were generated for two individual studies to test *H*_0*i*_: *μ*_*i*_ *≤* 0 *vs. H*_1*i*_: *μ*_1*i*_ *>* 0; *i* = 1, 2. The sample sizes for each study were generated from a Poisson distribution with a mean of 50. The standard deviations *σ*_*i*_ for each study were generated from the Uniform(2,4) distribution. Statistics under the individual null hypothesis were generated using *μ* = 0 (panel a) and *μ* = − 2 (panel b). The value of N.alt is the number of test statistics arising from parameters that lie in the alternative parameter space, with values of the mean given on the x-axis.

Figure 2 shows the power comparison between the proposed test and several other tests combining twenty individual studies under scenario 3. Clearly, the proposed test performs better than the other tests when only a small fraction of the individual tests are expected to be significant. In particular, when up to three of the twenty individual tests are drawn from the alternative space, the proposed test has considerably higher power. All the tests perform similarly when the number of significant individual tests increases. For instance, Figure 2 shows that when five of twenty individual test statistics are drawn from the alternative parameter space, all tests perform similarly with high power to reject the null hypothesis that none of the individual nulls is true.

**Figure 2:**
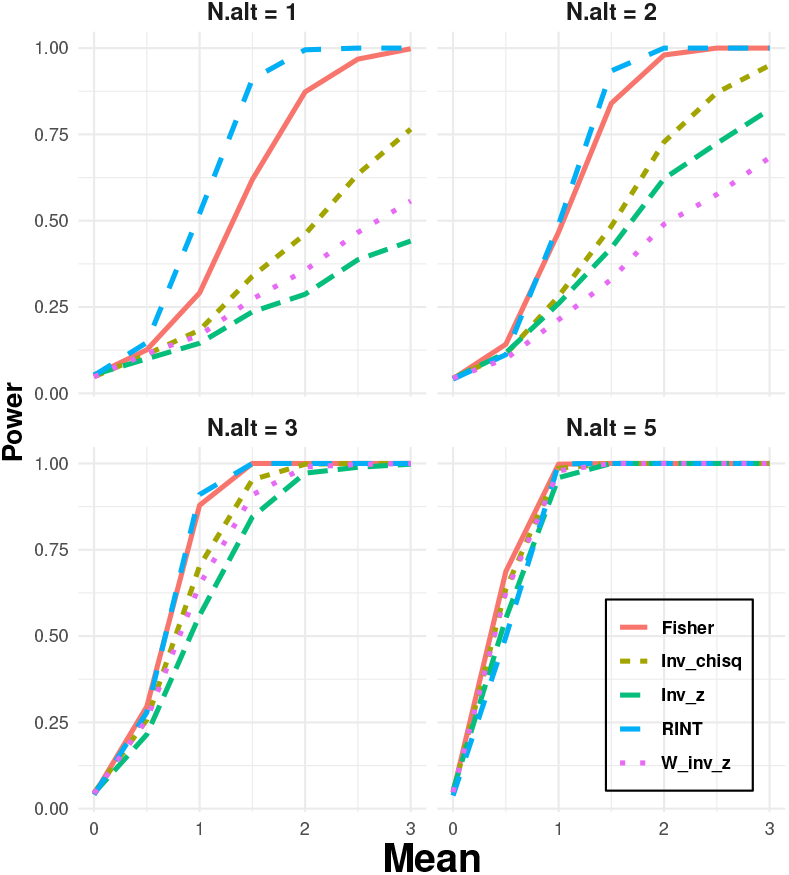
Plots of the power of the combination tests for data sets for which the P-values were generated under scenario 4 with *α* = 0.05. The x-axis of the plot shows the value of *μ* under the alternative space used to generate the individual test statistic, and the y-axis shows the power of several combination tests. P-values were generated for twenty individual studies to test *H*_0*i*_: *μ*_*i*_ *≤* 0 *vs. H*_1*i*_: *μ*_1*i*_ *>* 0; *i* = 1, 2, *· · ·*, 20. The sample sizes for each study were generated from a Poisson distribution with a mean of 50. The standard deviations *σ*_*i*_ for each study were generated from the Uniform(2,4) distribution. Statistics under the individual null hypothesis were generated using *μ* = 0. The value of N.alt is the number of test statistics arising from parameters that lie in the alternative parameter space, with values of the mean given on the x-axis.

Figure 3 shows the power comparison under scenario 10, where we conducted 100 individual tests, with 50 tests using a value of -2 for *μ* to generate the test statistic and the remainder using either 0 or a value greater than 0 for *μ*. As a result, we anticipate that 50 of the individual P-values will be close to 1. Figure 3 illustrates that in this scenario, the proposed test exhibits significantly higher power than the other tests, particularly when only a small number of individual tests are anticipated to be significant. Notably, the proposed test displays considerable power in rejecting the global null hypothesis, even if only one of 100 tests is expected to be significant. Conversely, other tests are powerless to reject the incorrect null hypothesis when some P-values are expected to be close to 1 and only a few P-values are expected to be smaller than the significance level. In this scenario, even when 30 of the 100 individual tests are expected to be significant, the proposed test outperforms all other tests.

**Figure 3:**
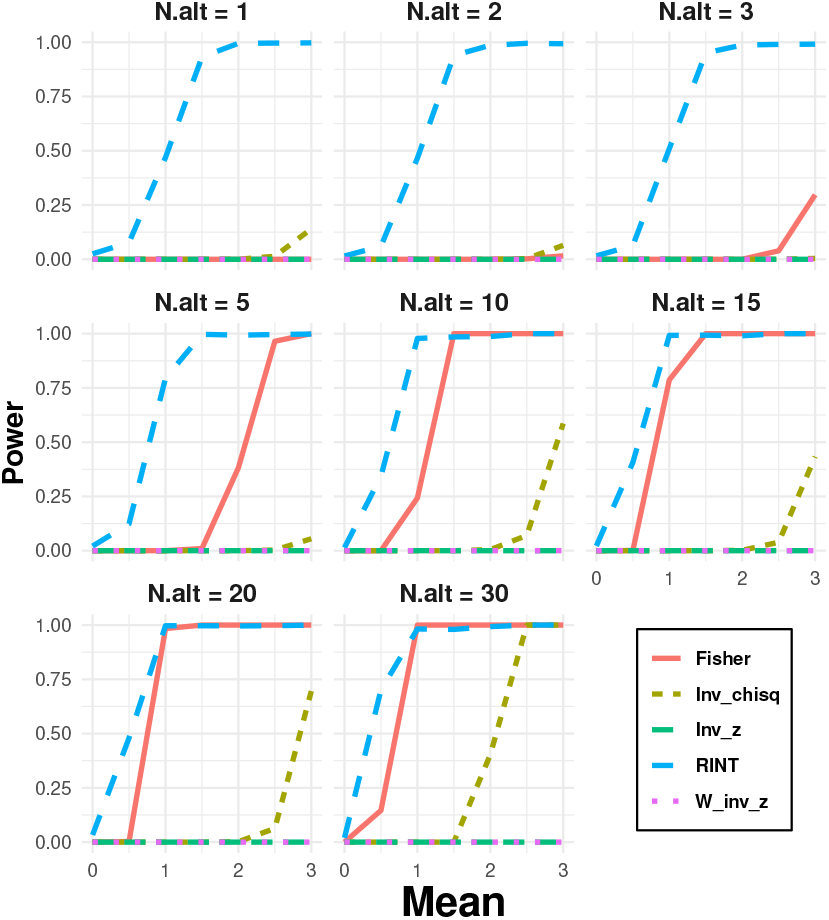
Plots of the power of the combination tests for data sets for which the P-values were generated under scenario 10 with *α* = 0.05. The x-axis of the plot shows the value of *μ* under the alternative space used to generate the individual test statistic, and the y-axis shows the power of several combination tests. P-values were generated for one hundred individual studies to test *H*_0*i*_: *μ*_*i*_ *≤* 0 *vs. H*_1*i*_: *μ*_1*i*_ *>* 0; *i* = 1, 2, *· · ·*, 100. The sample sizes for each study were generated from a Poisson distribution with a mean of 50. The standard deviations *σ*_*i*_ for each study were generated from the Uniform(2,4) distribution. Fifty statistics under the individual null hypothesis were generated using *μ* = − 2, making the corresponding P-values under the null hypothesis close to 1. The rest of the statistics under the null hypothesis were generated using *μ* = 0. The value of N.alt is the number of test statistics arising from parameters that lie in the alternative parameter space, with values of the mean given on the x-axis.

Figure 4 depicts the results of scenario 11, where we generated 20 individual test statistics for testing *μ*_*i*_ = 0 vs. *μ*_*i*_ ≠ 0; *i* = 1, 2, · · ·, 20. From this scenario, we observe similar results as in the other scenarios. The proposed test has higher power when a smaller number of individual tests are expected to be significant. The power of the other tests is similar when the number of significant individual tests increases.

**Figure 4:**
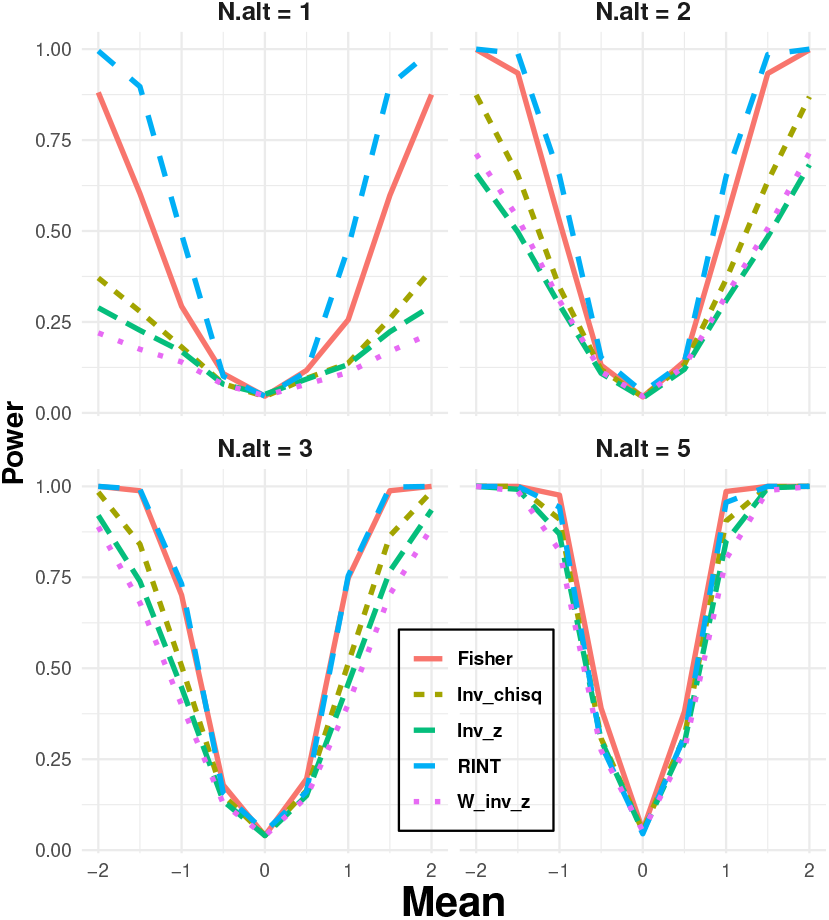
Plots of the power of the combination tests for data sets for which the P-values were generated under scenario 11 with *α* = 0.05. The x-axis of the plot shows the value of *μ* under the alternative space used to generate the individual test statistic, and the y-axis shows the power of several combination tests. P-values were generated for twenty individual studies to test *H*_0*i*_: *μ*_*i*_ = 0 *vs. H*_1*i*_: *μ*_1*i*_ ≠ 0; *i* = 1, 2, *· · ·*, 20. The sample sizes for each study were generated from a Poisson distribution with a mean of 50. The standard deviations *σ*_*i*_ for each study were generated from the Uniform(2,4) distribution. Statistics under the individual null hypothesis were generated using *μ* = 0. The value of N.alt is the number of test statistics arising from parameters that lie in the alternative parameter space, with values of the mean given on the x-axis.

The simulation study results indicate that the proposed test outperforms all of the traditional tests in scenarios where the number of test statistics from the alternative parameter space is much smaller than from the null parameter space. Specifically, the proposed test is more effective when combining P-values in cases for which only a fraction of the individual P-values are expected to be below the significance level. The simulation study’s findings for both significance level options (*α* = 0.05 and *α* = 0.01) are consistent with those discussed earlier (refer to supplementary materials at: https://drive.google.com/file/d/17QG94NIjrrLEG4KkikjKLLTdgU9NL5k9/view?usp=sharing). The proposed test performed better than all traditional tests, especially when only a portion of individual tests are expected to be significant or when some P-values are expected to be close to 1. It is noteworthy that all tests, including the proposed one, maintain the correct type I error for both significance-level options (refer to Appendix).

### 4.2 Application to Empirical Data

#### 4.2.1 Hybrid detection: Sistrurus rattlesnakes

We considered a data set consisting of two species of rattlesnakes (*Sistrurus catenatus* and *S. miliarius*) and two outgroup species (*Agkistrodon contortrix* and *A. piscivorus*) (Kubatko and Chifman, 2019). The species *S. catenatus* consists of subspecies *S. c. catenatus* (Sca), *S. c. edwardsii* (Sced), and *S. c. tergeminus* (Scter), and the species *Sistrurus miliarius* consists of subspecies *S. m. miliarius* (Smm), *S. m. barbouri* (Smb), and *S. m. streckeri* (Sms). The data set consists of aligned DNA sequences from 19 genes taken from 26 individual rattlesnakes. The species *S. catenatus* consists of 18 individuals (Sca: 9 individuals, Sced: 4 individuals, Scter: 5 individuals), and the species *S. miliarius* consists of 6 individuals (Smm: 1 individual, Smb: 3 individuals, and Sms: 2 individuals). The other 2 individuals are outgroups. Thus, the data set has DNA sequences on six distinct rattlesnake sub-species and two outgroups totaling 26 individual snakes. Since each DNA sequence is 8,466 base pairs in length, the sample size is 8,466 for all of the individual tests.

Several researchers postulated that a population of rattlesnakes in the *S. c. tergeminus* subspecies are hybrids because they have morphological characteristics that are intermediate between *S. c. catenatus* and other members of *S. c. tergeminus* and because these individuals are found in Missouri, on the edge of the range of *S. c. catenatus*. However, previous studies for the species considered here have found no evidence of hybridization (Gerard et al., 2011; Gibbs et al., 2011). In this study, we want to test the global null hypothesis that there are no hybrid species in the rattlesnake data vs. the alternative that at least one hybrid species is present. To perform this test, we will first compute the individual P-values testing each rattlesnake species for evidence of hybridization using the method proposed by Kubatko and Chifman (2019). Table 1 presents the P-values for all of these individual tests. Each row of Table 1 presents the P-value for testing whether the species in the middle is a hybrid of the two parents in the first and third columns. For instance, row 1 of Table 1 presents the P-value for testing whether Sced is a hybrid of parents Sca and Scter.

**Table 1:**
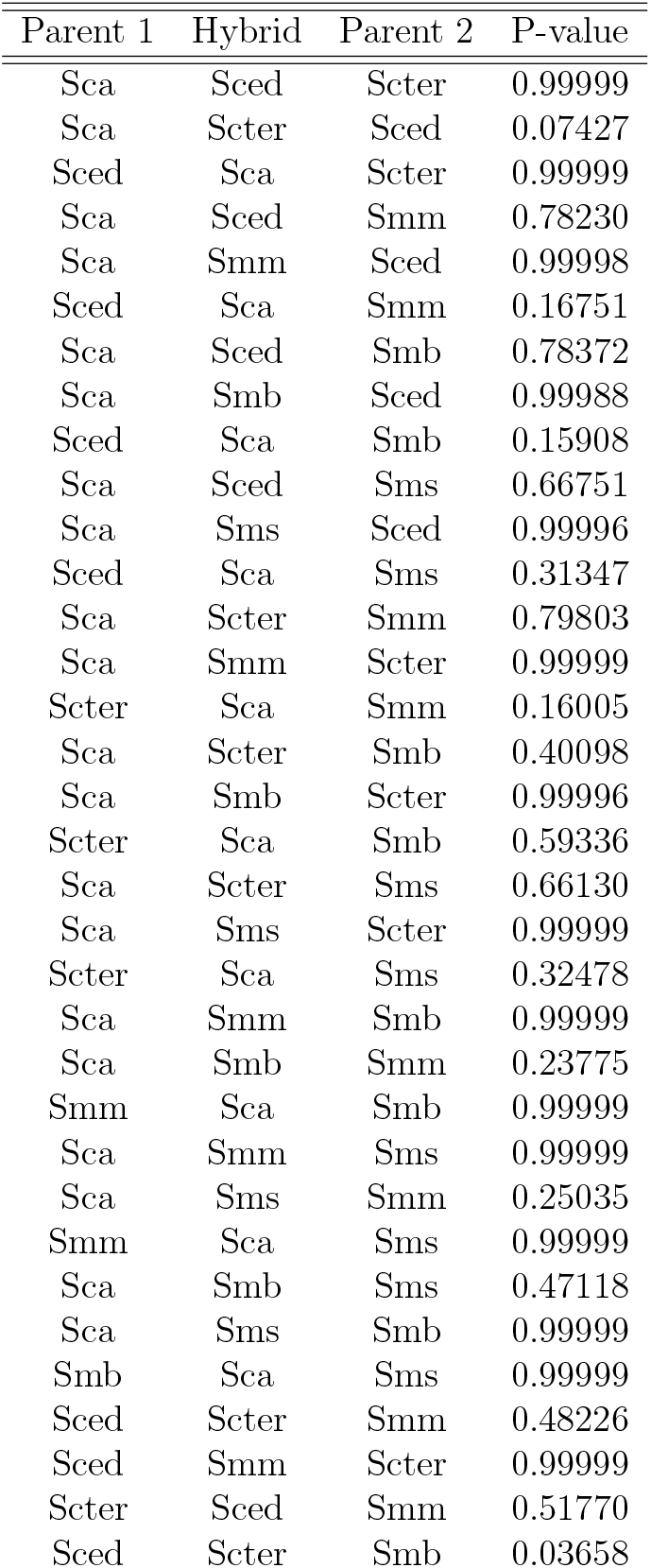
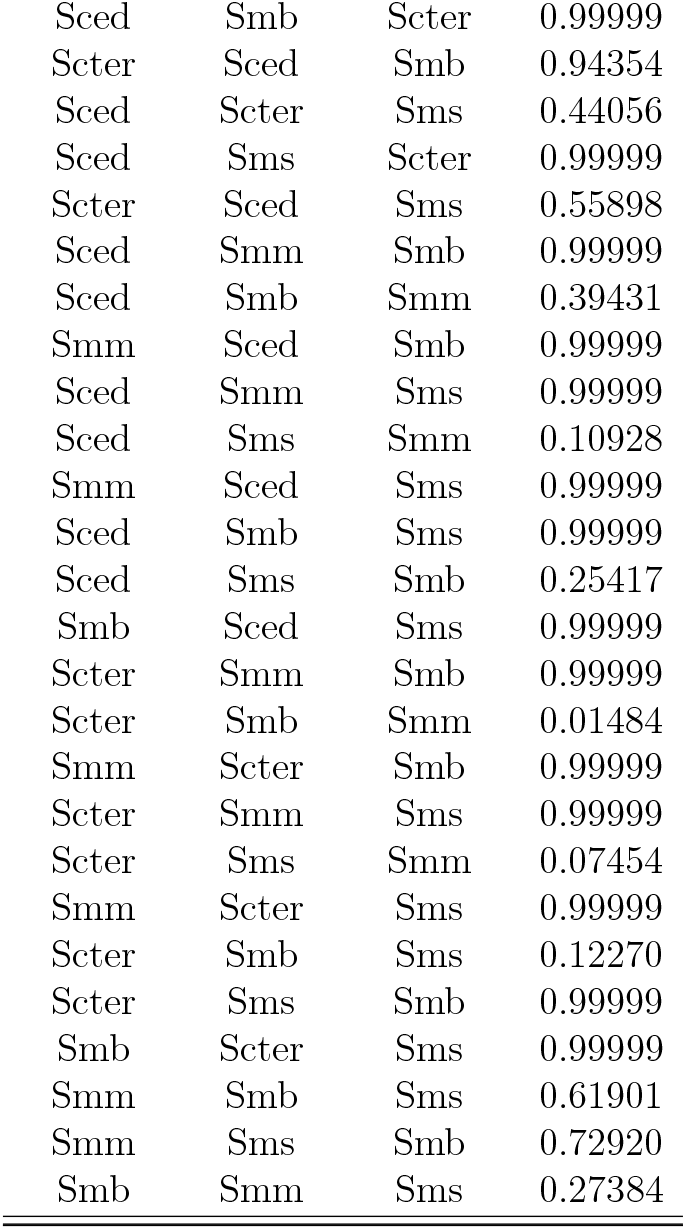
P-values for testing whether a particular rattlesnake species is a hybrid of two given parents. Each row represents an individual test. For example, row 1 represents the individual test of whether Sced is a hybrid of parents Sca and Scter.

| Parent 1 | Hybrid | Parent 2 | P-value |
| --- | --- | --- | --- |
| Sca | Sced | Scter | 0.99999 |
| Sca | Scter | Sced | 0.07427 |
| Sced | Sca | Scter | 0.99999 |
| Sca | Sced | Smm | 0.78230 |
| Sca | Smm | Sced | 0.99998 |
| Sced | Sca | Smm | 0.16751 |
| Sca | Sced | Smb | 0.78372 |
| Sca | Smb | Sced | 0.99988 |
| Sced | Sca | Smb | 0.15908 |
| Sca | Sced | Sms | 0.66751 |
| Sca | Sms | Sced | 0.99996 |
| Sced | Sca | Sms | 0.31347 |
| Sca | Scter | Smm | 0.79803 |
| Sca | Smm | Scter | 0.99999 |
| Scter | Sca | Smm | 0.16005 |
| Sca | Scter | Smb | 0.40098 |
| Sca | Smb | Scter | 0.99996 |
| Scter | Sca | Smb | 0.59336 |
| Sca | Scter | Sms | 0.66130 |
| Sca | Sms | Scter | 0.99999 |
| Scter | Sca | Sms | 0.32478 |
| Sca | Smm | Smb | 0.99999 |
| Sca | Smb | Smm | 0.23775 |
| Smm | Sca | Smb | 0.99999 |
| Sca | Smm | Sms | 0.99999 |
| Sca | Sms | Smm | 0.25035 |
| Smm | Sca | Sms | 0.99999 |
| Sca | Smb | Sms | 0.47118 |
| Sca | Sms | Smb | 0.99999 |
| Smb | Sca | Sms | 0.99999 |
| Sced | Scter | Smm | 0.48226 |
| Sced | Smm | Scter | 0.99999 |
| Scter | Sced | Smm | 0.51770 |
| Sced | Scter | Smb | 0.03658 |
| Sced | Smb | Scter | 0.99999 |
| Scter | Sced | Smb | 0.94354 |
| Sced | Scter | Sms | 0.44056 |
| Sced | Sms | Scter | 0.99999 |
| Scter | Sced | Sms | 0.55898 |
| Sced | Smm | Smb | 0.99999 |
| Sced | Smb | Smm | 0.39431 |
| Smm | Sced | Smb | 0.99999 |
| Sced | Smm | Sms | 0.99999 |
| Sced | Sms | Smm | 0.10928 |
| Smm | Sced | Sms | 0.99999 |
| Sced | Smb | Sms | 0.99999 |
| Sced | Sms | Smb | 0.25417 |
| Smb | Sced | Sms | 0.99999 |
| Scter | Smm | Smb | 0.99999 |
| Scter | Smb | Smm | 0.01484 |
| Smm | Scter | Smb | 0.99999 |
| Scter | Smm | Sms | 0.99999 |
| Scter | Sms | Smm | 0.07454 |
| Smm | Scter | Sms | 0.99999 |
| Scter | Smb | Sms | 0.12270 |
| Scter | Sms | Smb | 0.99999 |
| Smb | Scter | Sms | 0.99999 |
| Smm | Smb | Sms | 0.61901 |
| Smm | Sms | Smb | 0.72920 |
| Smb | Smm | Sms | 0.27384 |

To test the global null hypothesis that no hybrid species are present, Fisher’s test, the Inverse Chi-squared test, the Inverse Normal test, the Weighted Inverse Normal test, and the proposed test yielded P-values of 0.9991757, 1.0000000, 1.0000000, 1.0000000, and 0.8435300, respectively. The P-values obtained from the statistical tests are all quite large, indicating that the global null hypothesis of no hybrid species in the rattlesnake data set cannot be rejected. As a result, there is no evidence to suggest the presence of hybrid species in the data set. This result is in accordance with previous studies (Gerard et al., 2011; Gibbs et al., 2011).

#### 4.2.2 Hybrid detection: Yeast

We consider a subset of the data of Rokas et al. (2003) consisting of aligned DNA sequences for five yeast species: *Saccharomyces mikatae* (Smik), *S. kudriavzevii* (Skud), *S. bayanus* (Sbay), *S. kluyveri* (Sklu), and the outgroup fungus *Candida albicans* (Calb). Yu et al. (2011) found evidence of hybridization for these species. In particular, they found that the yeast species *S. kudriavzevii* (Skud) is a hybrid of parents *S. mikatae* (Smik) and *S. bayanus* (Sbay). The data consists of 120762 base pairs. Hence, the sample size *n* for each of the individual tests shown in Table 2 is 120762.

**Table 2:**
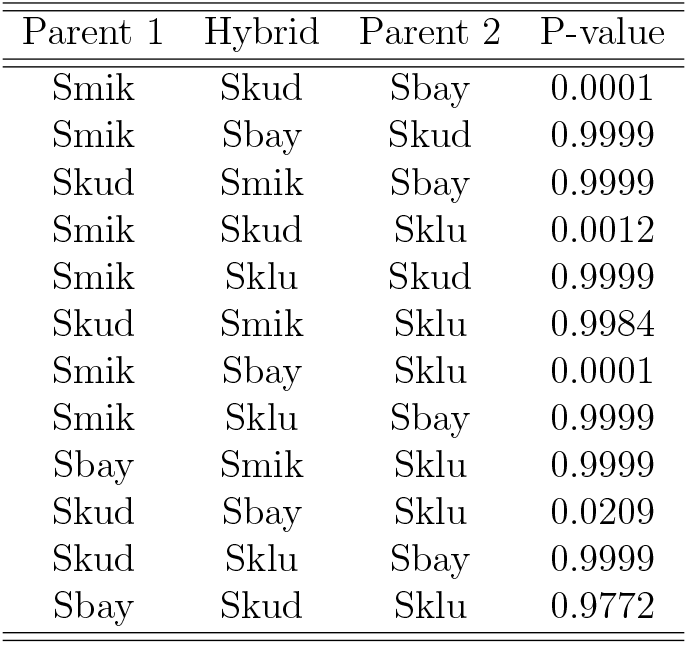
P-values for testing whether a particular yeast species is a hybrid of two given parents. Each row represents an individual test. For instance, the first row shows the result of testing whether the yeast species Skud is a hybrid of parents Smik and Sbay.

We test the global null hypothesis that none of the species in the yeast data set is hybrid against the alternative that at least one species is hybrid. To perform the global test, we conducted all possible individual tests considering each of the species in the yeast data set as a hybrid with two parents using the method proposed by Kubatko and Chifman (2019). The resulting P-values of all possible individual tests are presented in Table 2, where each row represents an individual test. For instance, the first row of Table 2 shows the result of testing whether the yeast species Skud is a hybrid of parents Smik and Sbay.

We use the proposed test along with several other existing combination tests to detect whether or not there are any hybrid species in the yeast data set. The resulting P-values from Fisher’s test, the Inverse Chi-squared test, the Inverse Normal test, the Weighted Inverse Normal test, and the proposed test are 0.0001204994, 0.9999877152, 0.9999897732, 0.9999897732, 0.0005800000, respectively. The P-values obtained from all of the statistical tests except for the proposed test and Fisher’s combination test are all quite large, indicating that the global null hypothesis of no hybrid species in the yeast data set cannot be rejected by any of the tests other than the proposed test and Fisher’s combination test. This result is in accordance with the findings of other researchers in the literature (Haque and Kubatko, 2023; Yu et al., 2011).

#### 4.2.3 Patient Rehabilitation Study

The data set is taken from Bachmann et al. (2010) and Riley et al. (2011). These studies conducted a meta-analysis of 12 randomized trials comparing patients receiving geriatric care versus patients receiving standard care on the effect of patient rehabilitation. The resulting odds ratios (ORs) and confidence intervals (CIs) from the 12 randomized trials are shown in Table 3.

**Table 3:** Odds ratios (ORs), 95% confidence intervals (CI), and P-values from the 12 randomized trials taken from Bachmann et al. (2010) and Riley et al. (2011).

| Study | OR | 95% CI | P-value |
| --- | --- | --- | --- |
| 1 | 1.11 | (0.51, 2.49) | 0.79970 |
| 2 | 0.97 | (0.78, 1.21) | 0.79422 |
| 3 | 1.13 | (0.73, 1.72) | 0.59899 |
| 4 | 1.08 | (0.42, 2.75) | 0.87931 |
| 5 | 0.88 | (0.39, 1.95) | 0.73626 |
| 6 | 1.28 | (0.71, 2.30) | 0.40873 |
| 7 | 1.19 | (0.69, 2.08) | 0.51677 |
| 8 | 3.82 | (1.37, 10.60) | 0.00962 |
| 9 | 1.06 | (0.63, 1.79) | 0.81974 |
| 10 | 2.95 | (1.54, 5.63) | 0.00097 |
| 11 | 2.36 | (1.18, 4.72) | 0.01417 |
| 12 | 1.68 | (1.05, 2.70) | 0.02892 |

**Table 4:** Type I error rate from the proposed method and all the other compared methods under all scenarios when the level of significance (*α*) is 0.05.

| Method | Power | Scenario | Significance level ($\alpha$) |
| --- | --- | --- | --- |
| Fisher | 0.0517 | Scenario 1 | 0.0500 |
| Inv_chisq | 0.0514 | Scenario 1 | 0.0500 |
| Inv_z | 0.0507 | Scenario 1 | 0.0500 |
| W_inv_z | 0.0509 | Scenario 1 | 0.0500 |
| New_z | 0.0519 | Scenario 1 | 0.0500 |
| Fisher | 0.0000 | Scenario 2 | 0.0500 |
| Inv_chisq | 0.0000 | Scenario 2 | 0.0500 |
| Inv_z | 0.0000 | Scenario 2 | 0.0500 |
| W_inv_z | 0.0000 | Scenario 2 | 0.0500 |
| New_z | 0.0000 | Scenario 2 | 0.0500 |
| Fisher | 0.0000 | Scenario 3 | 0.0500 |
| Inv_chisq | 0.0000 | Scenario 3 | 0.0500 |
| Inv_z | 0.0000 | Scenario 3 | 0.0500 |
| W_inv_z | 0.0000 | Scenario 3 | 0.0500 |
| New_z | 0.0000 | Scenario 3 | 0.0500 |
| Fisher | 0.0544 | Scenario 4 | 0.0500 |
| Inv_chisq | 0.0523 | Scenario 4 | 0.0500 |
| Inv_z | 0.0530 | Scenario 4 | 0.0500 |
| W_inv_z | 0.0534 | Scenario 4 | 0.0500 |
| New_z | 0.0501 | Scenario 4 | 0.0500 |
| Fisher | 0.0517 | Scenario 5 | 0.0500 |
| Inv_chisq | 0.0494 | Scenario 5 | 0.0500 |
| Inv_Z | 0.0500 | Scenario 5 | 0.0500 |
| W_inv_Z | 0.0477 | Scenario 5 | 0.0500 |
| New_Z | 0.0484 | Scenario 5 | 0.0500 |
| Fisher | 0.0499 | Scenario 6 | 0.0500 |
| Inv_chisq | 0.0519 | Scenario 6 | 0.0500 |
| Inv_Z | 0.0519 | Scenario 6 | 0.0500 |
| W_inv_Z | 0.0521 | Scenario 6 | 0.0500 |
| New_Z | 0.0511 | Scenario 6 | 0.0500 |
| Fisher | 0.0447 | Scenario 7 | 0.0500 |
| Inv_chisq | 0.0463 | Scenario 7 | 0.0500 |
| Inv_Z | 0.0456 | Scenario 7 | 0.0500 |
| W_inv_Z | 0.0470 | Scenario 7 | 0.0500 |
| New_Z | 0.0486 | Scenario 7 | 0.0500 |
| Fisher | 0.0000 | Scenario 8 | 0.0500 |
| Inv_chisq | 0.0000 | Scenario 8 | 0.0500 |
| Inv_Z | 0.0000 | Scenario 8 | 0.0500 |
| W_inv_Z | 0.0000 | Scenario 8 | 0.0500 |
| New_Z | 0.0000 | Scenario 8 | 0.0500 |
| Fisher | 0.0481 | Scenario 9 | 0.0500 |
| Inv_chisq | 0.0489 | Scenario 9 | 0.0500 |
| Inv_Z | 0.0490 | Scenario 9 | 0.0500 |
| W_inv_Z | 0.0490 | Scenario 9 | 0.0500 |
| New_Z | 0.0490 | Scenario 9 | 0.0500 |
| Fisher | 0.0000 | Scenario 10 | 0.0500 |
| Inv_chisq | 0.0000 | Scenario 10 | 0.0500 |
| Inv_Z | 0.0000 | Scenario 10 | 0.0500 |
| W_inv_Z | 0.0000 | Scenario 10 | 0.0500 |
| New_Z | 0.0204 | Scenario 10 | 0.0500 |
| Fisher | 0.0473 | Scenario 11 | 0.0500 |
| Inv_chisq | 0.0502 | Scenario 11 | 0.0500 |
| Inv_Z | 0.0482 | Scenario 11 | 0.0500 |
| W_inv_Z | 0.0504 | Scenario 11 | 0.0500 |
| New_Z | 0.0493 | Scenario 11 | 0.0500 |

**Table 5:** Type I error rate from the proposed method and all the other compared methods under all scenarios when the level of significance (*α*) is 0.01.

| Method | Power | Scenario | Significance level ($\alpha$) |
| --- | --- | --- | --- |
| Fisher | 0.0096 | Scenario 1 | 0.0100 |
| Inv_chisq | 0.0084 | Scenario 1 | 0.0100 |
| Inv_z | 0.0086 | Scenario 1 | 0.0100 |
| W_inv_z | 0.0083 | Scenario 1 | 0.0100 |
| New_z | 0.0100 | Scenario 1 | 0.0100 |
| Fisher | 0.0000 | Scenario 2 | 0.0100 |
| Inv_chisq | 0.0000 | Scenario 2 | 0.0100 |
| Inv_z | 0.0000 | Scenario 2 | 0.0100 |
| W_inv_z | 0.0000 | Scenario 2 | 0.0100 |
| New_z | 0.0000 | Scenario 2 | 0.0100 |
| Fisher | 0.0000 | Scenario 3 | 0.0100 |
| Inv_chisq | 0.0000 | Scenario 3 | 0.0100 |
| Inv_z | 0.0000 | Scenario 3 | 0.0100 |
| W_inv_z | 0.0000 | Scenario 3 | 0.0100 |
| New_z | 0.0000 | Scenario 3 | 0.0100 |
| Fisher | 0.0087 | Scenario 4 | 0.0100 |
| Inv_chisq | 0.0107 | Scenario 4 | 0.0100 |
| Inv_z | 0.0109 | Scenario 4 | 0.0100 |
| W_inv_z | 0.0101 | Scenario 4 | 0.0100 |
| New_z | 0.0100 | Scenario 4 | 0.0100 |
| Fisher | 0.0096 | Scenario 5 | 0.0100 |
| Inv_chisq | 0.0109 | Scenario 5 | 0.0100 |
| Inv_z | 0.0104 | Scenario 5 | 0.0100 |
| W_inv_z | 0.0099 | Scenario 5 | 0.0100 |
| New_z | 0.0091 | Scenario 5 | 0.0100 |
| Fisher | 0.0107 | Scenario 6 | 0.0100 |
| Inv_chisq | 0.0106 | Scenario 6 | 0.0100 |
| Inv_z | 0.0104 | Scenario 6 | 0.0100 |
| W_inv_z | 0.0109 | Scenario 6 | 0.0100 |
| New_z | 0.0097 | Scenario 6 | 0.0100 |
| Fisher | 0.0096 | Scenario 7 | 0.0100 |
| Inv_chisq | 0.0109 | Scenario 7 | 0.0100 |
| Inv_z | 0.0110 | Scenario 7 | 0.0100 |
| W_inv_z | 0.0114 | Scenario 7 | 0.0100 |
| New_z | 0.0109 | Scenario 7 | 0.0100 |
| Fisher | 0.0000 | Scenario 8 | 0.0100 |
| Inv_chisq | 0.0000 | Scenario 8 | 0.0100 |
| Inv_z | 0.0000 | Scenario 8 | 0.0100 |
| W_inv_z | 0.0000 | Scenario 8 | 0.0100 |
| New_z | 0.0000 | Scenario 8 | 0.0100 |
| Fisher | 0.0096 | Scenario 9 | 0.0100 |
| Inv_chisq | 0.0094 | Scenario 9 | 0.0100 |
| Inv_z | 0.0096 | Scenario 9 | 0.0100 |
| W_inv_z | 0.0096 | Scenario 9 | 0.0100 |
| New_z | 0.0111 | Scenario 9 | 0.0100 |
| Fisher | 0.0000 | Scenario 10 | 0.0100 |
| Inv_chisq | 0.0000 | Scenario 10 | 0.0100 |
| Inv_z | 0.0000 | Scenario 10 | 0.0100 |
| W_inv_z | 0.0000 | Scenario 10 | 0.0100 |
| New_z | 0.0046 | Scenario 10 | 0.0100 |
| Fisher | 0.0101 | Scenario 11 | 0.0100 |
| Inv_chisq | 0.0118 | Scenario 11 | 0.0100 |
| Inv_z | 0.0118 | Scenario 11 | 0.0100 |
| W_inv_z | 0.0119 | Scenario 11 | 0.0100 |
| New_z | 0.0083 | Scenario 11 | 0.0100 |

Riley et al. (2011) performed a random effect meta-analysis to estimate the overall OR. However, the goodness of fit test showed inadequate fit for the random effect model (Chen et al., 2015). Hence, it might be reasonable to use a combination test instead to find the overall P-value. Using the method of Chen et al. (2014), we observe the P-values for the individual studies as 0.79970, 0.79422, 0.59899, 0.87931, 0.73626, 0.40873, 0.51677, 0.00962, 0.81974, 0.00097, 0.01417, and 0.02892. Fisher’s test, the Inverse Chi-squared test, the Inverse Normal test, the Weighted Inverse Normal test, and the proposed test yielded P-values of 0.005709405, 0.186228113, 0.070941510, 0.521506484, 0.019670000, respectively, when tested for the global null hypothesis that there is no association between the type of care received and patient rehabilitation.

All tests except Fisher’s test and the proposed test had large P-values, resulting in a failure to reject the null hypothesis that patient rehabilitation has no association with the type of patient care (geriatric care versus standard care). The results from the proposed test and Fisher’s combination test are consistent with findings reported in other studies (Bachmann et al., 2010; Chen, Yang, Liu, Yang, Li and Yang, 2014; Riley et al., 2011).

## 5 Discussion and Conclusion

We propose a combination test based on the normal distribution that can combine independent P-values to test a global null hypothesis. One of the key advantages of this test is its ability to identify false null hypotheses even when only a small fraction of the individual tests are expected to be significant. Additionally, our test is robust in scenarios where some P-values are expected to be close to 1 and can accurately reject the null hypothesis even if only one individual test is significant. Importantly, our proposed test maintains the correct type I error rate in a variety of different scenarios. The simulation studies under several scenarios show that the proposed test is appropriate and has the highest power when a small fraction of the individual tests are expected to be significant. The real data applications show that the proposed test is one of the most powerful among the already existing tests and produces results that are supported by the existing literature. Also, the proposed test is shown to be robust through the simulation study since it not only is powerful when a small fraction of individual P-values are significant but also powerful when a large fraction of P-values are significant.

In addition to maintaining the correct type I error rate, our test also exhibits high power in a variety of scenarios. In many cases, our test has greater power than traditional tests, and in other scenarios, its power is similar or very close, making it a robust combination test. An additional benefit of our proposed test is its ability to incorporate the sample sizes of the individual tests as weights. On one hand, a test such as Weighted Inverse Normal takes sample sizes of individual studies into account but fails to assign adaptive weights to the independent P-values, leading to reduced power when only a small fraction of P-values are expected to be significant or when there are P-values close to 1. On the other hand, tests such as Fisher’s and Tippet’s minimum P-value approach are more effective for smaller P-values but fail to take different sample sizes into account. However, our proposed test takes sample sizes and the actual magnitude of the individual P-values into account simultaneously and adaptively, making it a robust test and appropriate in many applications.

While the proposed test has several advantages, it is important to acknowledge that traditional tests have their own strengths in specific scenarios. For example, Fisher’s test may have slightly higher power than our proposed test when none of the P-values are expected to be close to 1, and a significant number of individual P-values are expected to be smaller than the significance level. Another potential limitation of our proposed test is that the null distribution is not explicitly defined. We currently use the Uniform distribution to generate samples from the null distribution and calculate the critical value. Future work should focus on deriving a null distribution for our proposed test to enable the generation of a more precise critical value. However, it should be noted that generating data from Uniform distribution and calculating the critical values is efficient and does not take a notable amount of time.

## 6 Appendix

All the figures for the simulation study can be found here: https://drive.google.com/file/d/17QG94NIjrrLEG4KkikjKLLTdgU9NL5k9/view?usp=sharing

## References

Bachmann, S., Finger, C., Huss, A., Egger, M., Stuck, A. E. and Clough-Gorr, K. M. (2010), ‘Inpatient rehabilitation specifically designed for geriatric patients: systematic review and meta-analysis of randomised controlled trials’, BMJ 340.

Birnbaum, A. (1954), ‘Combining independent tests of significance’, Journal of the American Statistical Association 49(267), 559–574.

Blischak, P. D., Chifman, J., Wolfe, A. D. and Kubatko, L. S. (2018), ‘Hyde: a python package for genome-scale hybridization detection’, Systematic Biology 67(5), 821–829.

Chen, Z., Huang, H. and Liu, Q. (2014), ‘Detecting differentially methylated loci for multiple treatments based on high-throughput methylation data’, BMC Bioinformatics 15, 1–7.

Chen, Z., Huang, H. and Ng, H. K. T. (2012), ‘Design and analysis of multiple diseases genome-wide association studies without controls’, Gene 510(1), 87–92.

Chen, Z., Huang, H. and Ng, H. K. T. (2014), ‘An improved robust association test for gwas with multiple diseases’, Statistics & Probability Letters 91, 153–161.

Chen, Z., Huang, H. and Ng, H. K. T. (2016), ‘Testing for association in case-control genome-wide association studies with shared controls’, Statistical Methods in Medical Research 25(2), 954–967.

Chen, Z. and Ng, H. K. T. (2012), ‘A robust method for testing association in genomewide association studies’, Human Heredity 73(1), 26–34.

Chen, Z., Yang, W., Liu, Q., Yang, J. Y., Li, J. and Yang, M. Q. (2014), ‘A new statistical approach to combining p-values using gamma distribution and its application to genome-wide association study’, BMC Bioinformatics 15, 1–7.

Chen, Z., Zhang, G. and Li, J. (2015), ‘Goodness-of-fit test for meta-analysis’, Scientific Reports 5(1), 1–8.

Fisher, R. A. (1992), Statistical methods for research workers, in ‘Breakthroughs in Statistics’, Springer, pp. 66–70.

Gao, J., Pan, Z., Jiao, Z., Li, F., Zhao, G., Wei, Q., Pan, F. and Evangelou, E. (2012), ‘Tph2 gene polymorphisms and major depression–a meta-analysis’, PloS ONE 7(5), e36721.

Gerard, D., Gibbs, H. L. and Kubatko, L. (2011), ‘Estimating hybridization in the presence of coalescence using phylogenetic intraspecific sampling’, BMC Evolutionary Biology 11(1), 1–12.

Gibbs, H. L., Murphy, M. and Chiucchi, J. E. (2011), ‘Genetic identity of endangered massasauga rattlesnakes (Sistrurus sp.) in Missouri’, Conservation Genetics 12, 433–439.

Good, I. (1955), ‘On the weighted combination of significance tests’, Journal of the Royal Statistical Society: Series B (Methodological) 17(2), 264–265.

Haque, M. R. and Kubatko, L. (2023), ‘A global test of hybrid ancestry from genomescale data’, bioRxiv. URL: https://www.biorxiv.org/content/early/2023/02/27/2023.02.24.529943

Kong, S. and Kubatko, L. S. (2021), ‘Comparative performance of popular methods for hybrid detection using genomic data’, Systematic Biology 70(5), 891–907.

Krishnamoorthy, K., Lv, S. and Murshed, M. M. (2022), ‘Combining independent tests for a common parameter of several continuous distributions: a new test and power comparisons’, Communications in Statistics-Simulation and Computation pp. 1–20.

Kubatko, L. S. and Chifman, J. (2019), ‘An invariants-based method for efficient identification of hybrid species from large-scale genomic data’, BMC Evolutionary Biology 19(1), 1–13.

Lancaster, H. (1961), ‘The combination of probabilities: an application of orthonormal functions’, Australian Journal of Statistics 3(1), 20–33.

Lipták, T. (1958), ‘On the combination of independent tests’, Magyar Tud Akad Mat Kutato Int Kozl 3, 171–197.

Liu, Y. and Xie, J. (2020), ‘Cauchy combination test: a powerful test with analytic p-value calculation under arbitrary dependency structures’, Journal of the American Statistical Association 115(529), 393–402.

Meng, C. and Kubatko, L. S. (2009), ‘Detecting hybrid speciation in the presence of incomplete lineage sorting using gene tree incongruence: a model’, Theoretical Population Biology 75(1), 35–45.

Mosteller, F. and Bush, R. R. (1954), Selected quantitative techniques, Addison-Wesley.

Pearson, E. S. (1938), ‘The probability integral transformation for testing goodness of fit and combining independent tests of significance’, Biometrika 30(1/2), 134–148.

Riley, R. D., Higgins, J. P. and Deeks, J. J. (2011), ‘Interpretation of random effects meta-analyses’, BMJ 342.

Rokas, A., Williams, B. L., King, N. and Carroll, S. B. (2003), ‘Genomescale approaches to resolving incongruence in molecular phylogenies’, Nature 425(6960), 798–804.

Stouffer, S. A., Suchman, E. A., DeVinney, L. C., Star, S. A. and Williams Jr, R. M. (1949), ‘The American soldier: Adjustment during army life (Studies in social psychology in World War II), vol. 1’.

Tippett, L. H. C. et al. (1931), ‘The methods of statistics.’, The Methods of Statistics..

Whitlock, M. C. (2005), ‘Combining probability from independent tests: the weighted z-method is superior to Fisher’s approach’, Journal of Evolutionary Biology 18(5), 1368–1373.

Wicke, K., Haque, M. R. and Kubatko, L. (2023), ‘Effects of phylogenetic variation on prioritization of species for conservation’, bioRxiv. URL: https://www.biorxiv.org/content/early/2023/01/21/2023.01.21.525012

Yu, Y., Than, C., Degnan, J. H. and Nakhleh, L. (2011), ‘Coalescent histories on phylogenetic networks and detection of hybridization despite incomplete lineage sorting’, Systematic Biology 60(2), 138–149.

